# Photoreceptor precursor cell integration into rodent retina after treatment with novel glycopeptide PKX-001

**DOI:** 10.1101/2020.11.22.393439

**Authors:** Ishaq A. Viringipurampeer, Anat Yanai, Cheryl Y. Gregory-Evans, Kevin Gregory-Evans

**Author notes:** **Corresponding Author:** Kevin Gregory-Evans MD, PhD.

## Abstract

Cell replacement therapy is emerging as an important approach in novel treatments for neurodegenerative diseases. Many problems remain, in particular improvements are needed in the survival of transplanted cells and increasing functional integration into host tissue. These problems arise because of immune rejection, suboptimal precursor cell type, trauma during cell transplantation, toxic compounds released by dying tissues and nutritional deficiencies. We recently developed an *ex vivo* system to facilitate identification of factors contributing to the death of transplanted neuronal (photoreceptor) and showed 2.8-fold improvement in transplant cell survival after pre-treatment with a novel glycopeptide (PKC-100). In this study we extended these studies to look at cell survival, maturation and functional integration in an *in vivo* rat model of rhodopsin-mutant retinitis pigmentosa causing blindness. We found that only when human photoreceptor precursor cells (PPCs) were pre-incubated with PKX-100 prior to transplantation, did the cells integrate and mature into cone photoreceptors expressing S-opsin or L/M opsin. In addition, ribbon synapses were observed in the transplanted cells suggesting they were making synaptic connections with the host tissue. Furthermore, optokinetic tracking and electroretinography responses *in vivo* were significantly improved compared to cell transplants without PKX-100 pre-treatment. These data demonstrate that PKX-100 promotes significant long-term stem cell survival *in vivo*, providing a platform for further investigation towards the clinical application to repair damaged or diseased retina.

## INTRODUCTION

Cell transplantation is emerging as an important innovation in future treatments of many neurological diseases such as stroke and the many subtypes of neurodegeneration (such as Alzheimer’s disease). (Bliss et al., 2007; Stone et al., 2013; Hao et al., 2014). A key feature of cell transplantation is the possibility of recovery of function in such diseases. Recently, photoreceptor precursor cells (PPCs) have been studied as possible treatments for retinal diseases such as age-related macular degeneration (AMD) and retinitis pigmentosa (RP) (Klassen, 2015).

An important issue in this work is the low survival rate of retinal precursor cells after transplantation *in vivo*. (Stone et al., 2013; Warre-Cornish et al., 2014). Many factors could be responsible for this including immune rejection, suboptimal cell types being used for transplantation, trauma during cell delivery, nutritional deficits in the target tissue and toxic compounds secreted from the target tissue (Ma et al., 2011; West et al., 2012; Singh et al., 2013).

Recently we highlighted the possible role of necrotic release of prostaglandins from degenerating host tissue (Ricciotti and FitzGerald, 2011) and the secondary triggering of inflammation (Zhou and Yuan, 2014) as possible contributory factors in limiting the lifespan of cells transplanted into retinal tissue (Yanai et al, 2015).

Previously we showed that PPC survival can be significantly improved when cells are pretreated with PKX-001 (previously known as anti-ageing glycoprotein AAGP) in an *ex-vivo* model of human retinal dystrophy (Yanai et al, 2015). In this study we extend the work into studies of cell survival, maturation and functional integration into rodent retina exhibiting a retinal dystrophy due to rhodopsin gene mutation (SD-*Foxn1* Tg(*S334ter*)3Lav rat).

## MATERIALS AND METHODS

### Preparation of PPCs

Peripheral blood was collected from a healthy volunteer into a BD Vacutainer tube and mononuclear cells were isolated using the SepMate kit (STEMCELL Technologies) according to the manufacturer’s protocol. The purified cells were re-suspended in fetal bovine serum with DMSO (9:1) and stored at -80°C. The cells were reprogramed into iPSCs by the Centre for Commercialization of Regenerative Medicine (CCRM; Toronto). Photoreceptor precursor cells (PPCs) were derived from human iPSC as per the protocol by Yanai et al., 2013 and Yanai et al., 2015. To confirm PPC identity and functionality, immunocytochemistry for photoreceptor markers S-Opsin, L/M-Opsin, Recoverin and CRX was performed. In addition, gene expression of pluripotency (Nanog and SOX2) and photoreceptors (CRX, NRL, Opsin, Recoverin and RHO) markers was analyzed by qPCR as previously described (Yanai et al., 2013; Yanai et al., 2015) using an iScript cDNA synthesis kit (Bio-Rad) and TaqMan Primer/probe sets for each gene. Gene expression was determined using a ViiA™ Real Time PCR System (Applied Biosystems). Data were analyzed by the comparative C_T_ method (Livak et al., 2001). Each reaction was undertaken on three separate occasions, and for each of these, individual samples were diluted into three replicates (N=9).

### Treatment of PPCs with PKX-001

At 24 hrs prior to cell transplantation, PPCs were treated with 4 mg/mL of PKX-001 in culture media ((DMEM/F12, 2% B-27 and 1% N-2 supplements,1% sodium pyruvate and 1% Non-essential amino acids). The control group of cells received the same culture medium without PKX-001. Cell culture plates were incubated at 37°C with 5% CO_2_ and a humidified environment.

### Labeling of PPCs

Following a 24hr incubation with or without PKX-001, PPCs were labeled with Cell Trace Far Red DDAO-SE according to manufacturer specifications (Thermo Fisher Scientific). After labeling, cells were dissociated by adding 1 mL of TrypLE™ Express Enzyme (Thermo Fisher Scientific) to each well for 3 min, after which culture media was added back to the wells to neutralize trypsin. Cells were collected, centrifuged for 5 min at 1500 rpm, supernatants were discarded and cells were re-suspended in sterile Dulbecco PBS (Thermo Fisher Scientific). Cell viability was determined by Trypan blue. Labeled cells were adjusted to a concentration of 10^6^ viable cells/mL and kept on ice prior to transplantation.

### Animal model

All animal procedures were carried out with approval of the Animal Care Committee of the University of British Columbia and in accordance with the Association for Research in Vision and Ophthalmology Statement for the Use of Animals in Ophthalmic and Vision Research.

Breeding males and females were generously provided by Dr. M.J. Seiler (Reeve-Irvine Research Center, University of California, Irvine, CA, USA). Heterozygous +/*Foxn1*rnu, S334ter transgene (Tg)-positive females were bred with homozygous (*Foxn1*rnu/*Foxn1*rnu), S334ter Tg-positive males (Seiler MJ. *et al*, 2014). As all newborn pups are homozygous (*Foxn1*rnu/*Foxn1*rnu) no genotyping was required [all homozygous (*Foxn1*rnu/*Foxn1*rnu), S334ter Tg-positive males are nude and blind]. Rats entered the study in multiple cohorts when they were 21 days old, with animals in each cohort being randomized between the study groups. In all experiments the right eye of each animal was injected with PPCs.

For PPC transplantation, SD-*Foxn1* Tg(*S334ter*)3Lav rats were anesthetized with isoflurane and host-eyes were anesthetized with proparicaine (Alcon, Fort Worth, TX, USA) preceding insertion of lid-retractors. Pupils were dilated with 1% tropicamide (Bausch and Lomb, Rochester, NY, USA). Povidone-iodine (Alcon) antimicrobial solution was used to wash the eye, lids, surrounding skin, and fur of the recipient. Tear-gel (Alcon) was applied to the cornea throughout the surgery. Filter-sterilized 2% Nile Blue (Sigma-Aldrich, St Louis, MO, USA) dye (in dPBS) was used to mark limbal-conjunctiva at the temporal and inferior midlines for orientation of the eye. Host conjunctiva was then reflected (one-fourth–one-sixth circumference), followed by rotation of the globe by grasping the limbal-conjunctiva with micro-forceps to visualize the inferior-temporal sclera.

Approximately, 100,000 cells were aseptically injected into the sub-retinal space (100,000 cells in 10 *μ*L of sterile dPBS) using a 30-gauge needle attached to a 10 *μ*L Gastight Syringe (Hamilton, Reno, NV, USA). The needle was removed 1 min after the injection. Recipients were then allowed to recover in an incubator with host eyes treated with proparicaine analgesic drops, tobramycin (0.3%) ophthalmic antibiotic ointment, and tear gel (all from Alcon). No immune suppression was required as all the rats were nude, lacking a thymus-derived T-cell lymphocyte response.

### Electroretinography (ERG)

ERGs were recorded at 3, 4.5, and 6 months post-transplantation (3 rats per time point). To measure the electrical response of photoreceptors in the retina in response to a light stimulus, a full-field ERG test was performed. In brief, rats were dark-adapted overnight prior to ERG analysis, then anesthetized and placed on a heating pad to maintain a constant body temperature at 37 °C during the entire test. The cornea was anesthetized with topical 0.5% proparacaine hydrochloride (Bausch and Lomb, Rochester, NY, USA) and the pupils were dilated with 2.5% phenylephrine and 1% atropine. A drop of 2% hydroxy-propyl-methyl-cellulose was placed on each cornea to maintain corneal hydration. ERG responses of the retina to light flashes were recorded using an Espion E2 system with a Colordome mini-Ganzfeld stimulator (Diagnosys LLC, Lowell, MA, USA) by averaging 15 responses at a stimulus intensity of 3.16 c/d/s/m^2^. Light-adapted cone responses were carried out in 30c/d/m^2^ background light. The data were collected at 12 Hz flicker amplitude.

### Optokinetic tracking (OKT)

In order to examine visually driven behavioral responses in the rat, OKT (Douglas et al, 2015) was used at 3, 4.5, and 6 months post transplantation (3-5 rats per time point) to measure spatial frequency thresholds under photopic (light adapted) conditions (Bashar et al, 2016). A vertical sine wave grating (100% contrast) was projected as a virtual cylinder in three-dimensional space on computer monitors arranged in a quadrangle around a testing arena (OptoMotry; CerebralMechanics, Lethbridge, AB, Canada). Unrestrained rats were placed on the elevated platform at the center of the equipment. An observer used a video monitor to ensure that the virtual cylinder is kept centered on the animal’s head and records head movements in response to cylinder rotation. Visual acuity was expressed as cycles/degree (c/d).

### Immunostaining and confocal imaging

Three to five rats per group were sacrificed 3, 4.5, and 6-months post-transplantation and human PPC cell survival, maturation and integration were examined. Following euthanasia, transplant-recipient eyes were marked at the inferior midline and nasal equator with Blue dye (1% methylene blue) for orientation then enucleated and fixed in 4% paraformaldehyde for 24 h at 4°C. *<u>For flat mounts</u>*: neurosensory retina was obtained by blunt dissection from the eyecups, flattening the tissue onto Superfrost Plus microscope slides (Fisher), and mounting under a coverslip with DAPI Fluoromount-G (Southern Biotech; 1-2 drops). Nuclear staining was detected with excitation at 405 nm. Transplanted cells pre-labelled with Cell Trace Far Red DDAO-SE were visualized by excitation at 633nm. Flat mounts were examined under a 10X objective with a Zeiss LSM 800 super-resoution confocal microscope and initially assessed qualitatively by visual examination. To quantify PPCs, three 600µm^2^ z-stack maximum projection images for each treatment group were counted. Counting was masked to the treatment group to avoid observer bias. *<u>For immunohiostochemistry</u>*: The anterior segment of the eyes was removed and the retina *in situ* was cryopreserved in 30% sucrose overnight, snap frozen and embedded in Polyfreeze medium (Polysciences). Sagittal sections (∼ 16 µm thick) were incubated overnight at 4°C with primary antibody diluted in blocking buffer (2% normal goat serum, 0.1% Triton X-100 in PBS). After extensive washes in PBS-Tween 20, localization of antibody labeling was detected after a 1 hr incubation with secondary antibody diluted in PBS containing 2% normal goat serum and mounting under a coverslip with DAPI Fluoromount-G (Southern Biotech). PPCs were visualized by fluorescent microscopy and their numbers were assessed qualitatively by visual examination. The following primary antibodies were used: anti-CRX (in house preparation; Dilution 1:50, overnight incubation at 4°C), anti-STEM121 (Takara; Dilution 1:200, overnight incubation at 4°C), anti-Ribeye (clone16/CTBP2BD Transduction laboratories; Dilution 1:100, overnight incubation at 4°C), anti-S-opsin (Millipore; Dilution 1:300, overnight incubation at 4°C), anti-L/M-opsin (Millipore; Dilution 1:300, overnight incubation at 4°C). Secondary antibodies were either goat anti-rabbit or anti-mouse IgG (H+L) cross-adsorbed antibodies, Alexa Fluor 488 (both from Invitrogen, Dilution 1:200; 1 hr incubation at room temperature). Images were acquired using a Zeiss LSM800 super-resolution confocal microscope.

## RESULTS

### Characterization of PPCs

After differentiation of iPSCs into PPCs, their identity was confirmed by immunocytochemistry and qPCR prior to transplantation. By immunohistochemistry PPCs expressed CRX (an early photoreceptor marker), L/M-opsin and S-opsin (cone photoreceptor markers), and recoverin (late photoreceptor marker) (**Fig. 1A**). By qRT-PCR we found that in comparison to iPSCs, the PPCs expressed significantly higher levels of CRX, S-Opsin and Recoverin (**Fig. 1B**). In contrast, a lower level of pluripotency markers were expressed by PPCs than by iPSCs confirming the cells were in a differentiated state.

**Fig. 1.**
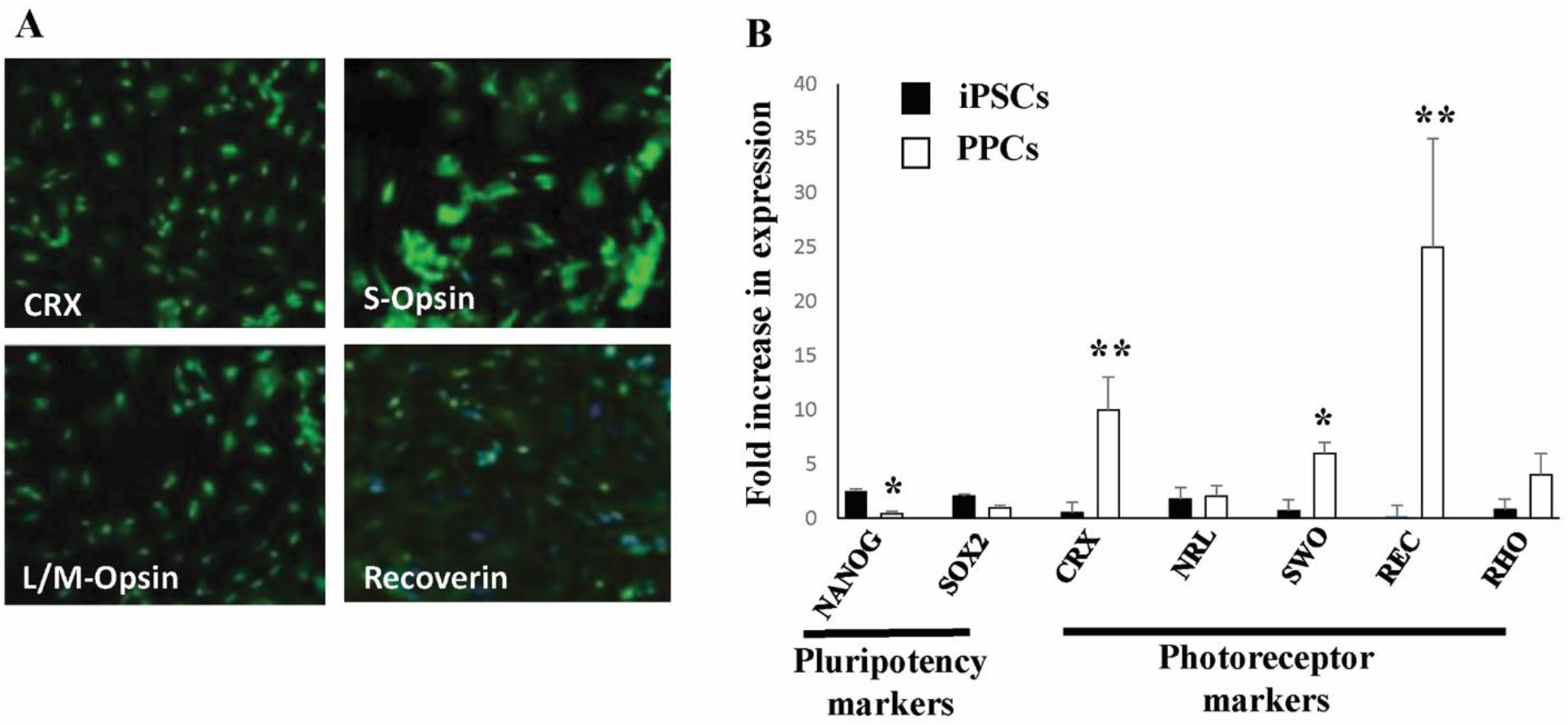
Characterization of PPCs. (**A**) Immunohistochemical labelling of PPCs with antibodies to CRX, S-Opsin, L/M-Opsin and Recoverin. (**B**) Quantification of real-time RT-PCR expression data for pluripotency markers (NANOG and SOX2) and photoreceptor markers (CRX, NRL, SW-opsin (SWO), Recoverin (REC), and rhodopsin (RHO) in iPSCs and in differentiated PPCs. Data presented as mean ± SEM. *P<0.001; **P<0.01 (N=9).

### Functional assessment of vision after transplantation

Approximately, 100,000 cells were aseptically injected into the sub-retinal space of the right eye in each animal. Prior to histological analysis at 3, 4.5 and 6 months, full field ERG testing was carried out to determine if there was a change in response of the retina to light in each treatment group. Photopic and scotopic waveforms decline rapidly within the first six months in the SD-*Foxn1* Tg(*S334ter*)3Lav rat model and only 12Hz flicker responses are retained in the control group at 6-months post-transplantation. We therefore reserved quantitative analysis for 12 Hz flicker responses (**Table 1**).

**Table 1.** 12Hz flicker data for each treatment group.

| Time | Mutant<br>3 Months |  | Mutant<br>4.5 Months |  | Mutant<br>6 Months |  | Wildtype<br>6 Months | Mutant<br>6 Months |
| --- | --- | --- | --- | --- | --- | --- | --- | --- |
| Treatment | Cells<br>only | Cells +<br>PKX-100 | Cells<br>only | Cells +<br>PKX-100 | Cells<br>only | Cells +<br>PKX-100 | None | None |
| Mean 12Hz<br>Flicker<br>Amplitude (μV) | 5.33<br>±<br>2.90<br>(N=3) | 14.3<br>±<br>1.86<br>(N=3) | 5.0<br>±<br>2.90<br>(N=3) | 15.0<br>±<br>2.90<br>(N=3) | 4.4<br>±<br>1.96<br>(N=5) | 12.2<br>±<br>2.35<br>(N=5) | 80.2<br>±<br>9.0<br>(N=5) | 4.3<br>±<br>2.1<br>(N=4) |
| *P* value:<br>cells versus<br>cells+PKX100 |  | 0.06 |  | 0.07 |  | *0.03 |  |  |

The mean value of 12 Hz flicker amplitude in the PKX-001-treated PPC group was higher than in the untreated PPC group at all time points but was only significant for the 6 month treatment group (P=0.03, N=5). Based on in-house historical data, the wildtype 12 Hz flicker amplitude in rats is approximately 80 μV. The values observed in the PKX-001-treated PPC group were in the 12-15μV range at 3, 4.5 and 6 months. Therefore, only a partial restoration of the ERG response was observed in this study.

To examine the behavioral response of the animals to light, we used optokinetic tracking (OKT) analysis. To standardize animal responses due to potentially different levels of natural degeneration between animals, the OKT response observed with the left (untreated) eye was subtracted from the response obtained with the right (treated) eye in the groups that received cell therapy with and without PKX-001 treatment. In the control group (no cell transplant), the response was normalized by subtracting the lower OKT response from the higher OKT response which can be right minus left eye or left minus right eye. This was done to avoid negative OKT values and to determine the absolute value for the response variation that can be expected from the eyes of the same animal.

At each time point, the animals transplanted with PPCs alone did not show any statistically significant improvement in OKT responses (**Fig. 2**). However, when cells were pretreated with PKX-001 prior to transplantation, the OKT response was significantly enhanced at 4.5 months and 6 months post-transplantation (P=0.05; N=3 and P=0.01; N=5, respectively). Healthy rat mean value of spatial frequency was 0.55 c/d (Douglas et al, 2015). The spatial frequency observed in the cell transplant group with PKX-001 treatment was in the range of 0.12–0.51 c/d at 3 months, 0.17–0.38 c/d at 4.5 months and 0.16–0.30 c/d at 6 months suggesting a significant retention of what in humans can be equated with visual acuity.

**Fig. 2.**
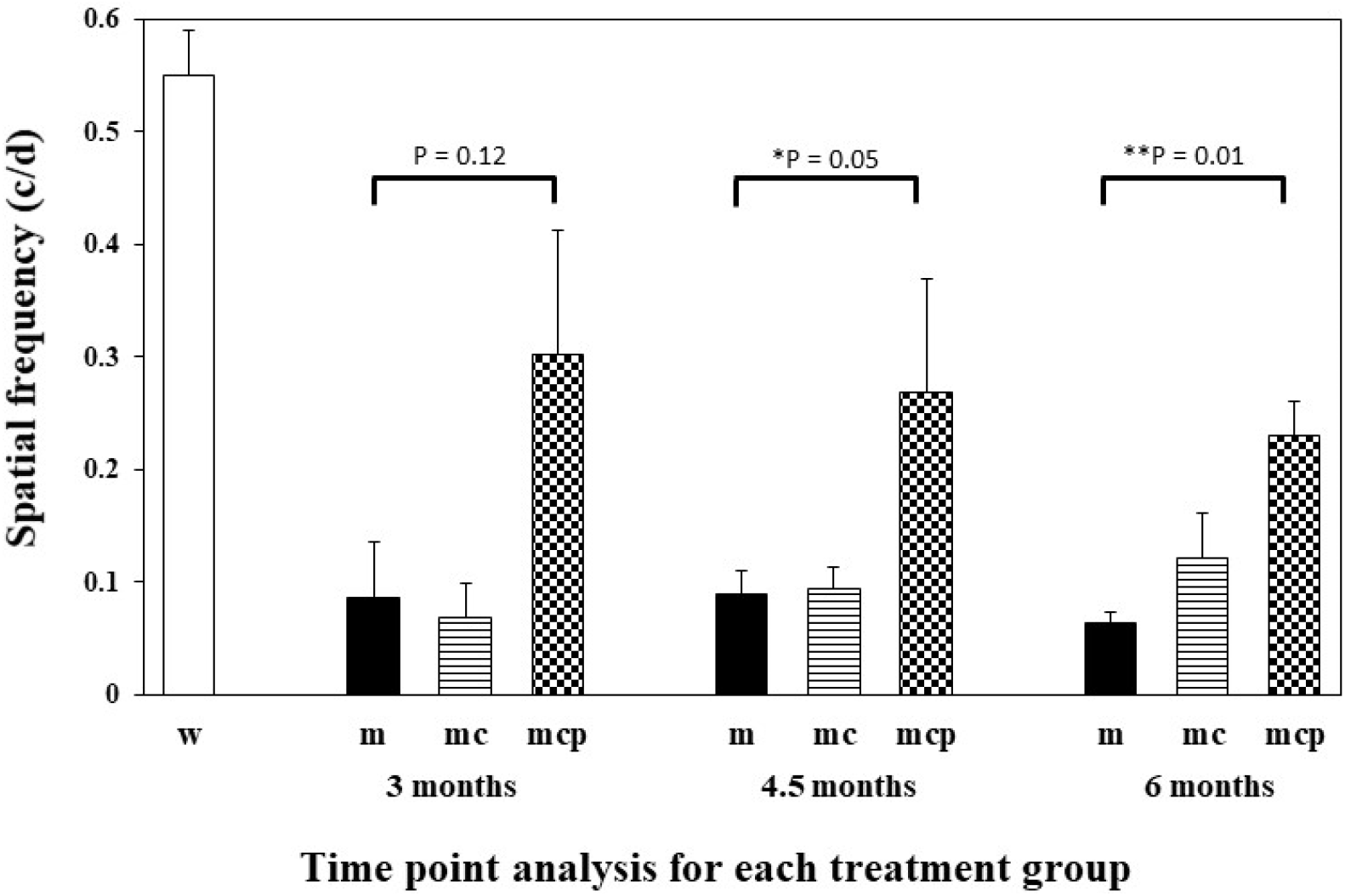
The effect of PKX-100 treatment on OKT tracking in rats receiving PPC transplantation. The comparisons were made between the following groups: untreated mutant rats (m); mutant rats receiving PPC transplants (mc) and mutant rats receiving PPC transplants that were pre-treated with PKX-100 (mcp). The spatial frequency of treated rats was compared to untreated animals at each time point. Data presented as mean ± SEM (Students *t* test; *P=0.05, N=3; **P=0.01, N=5). Wildtype (w) rat data presented for comparison.

### Survival of PPCs after transplantation

Flat mount retinas were collected from untreated control animals (no cell transplant), animals treated with PPCs without prior incubation with PKX-001 and animals treated with PPCs that had undergone prior PKX-001 incubation. Flat mounts were collected at 3 months, 4.5 month and six-month time points (**Fig. 3**). Qualitatively, there appeared to be more PPCs in the retinas when the PPCs had been pre-treated with PKX-100.

**Fig. 3.**
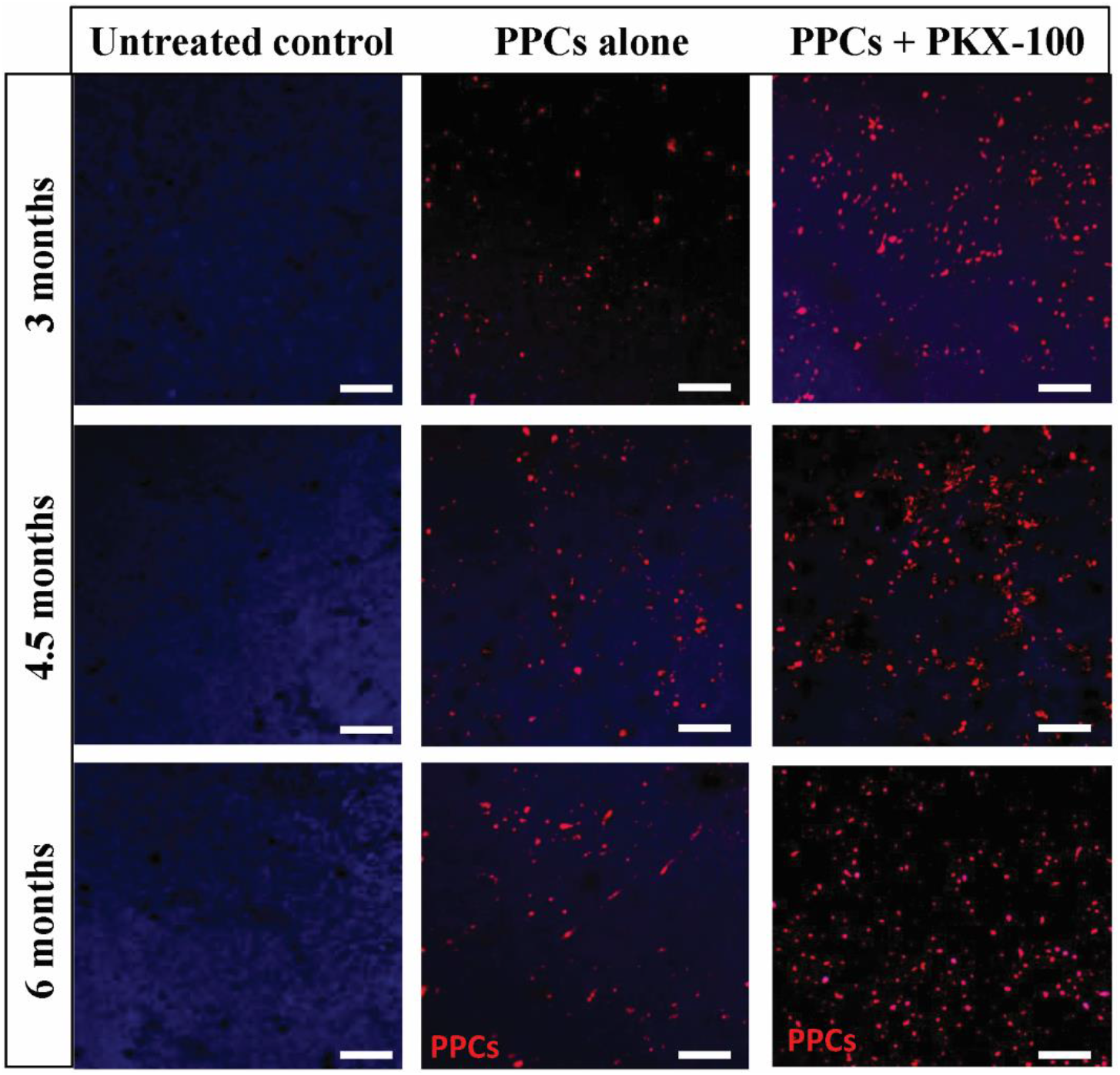
Representative confocal images of flat mount retinas from each treatment group at 3, 4.5 or 6 months post-transplantation. Photoreceptor precursor cells (PPCs) are labelled in red and retinal nuclei counterstained with DAPI. Size bar = 25µm.

To quantify the difference in cell counts between PPCs transplanted alone or pre-treated with PKX-100 prior to transplantation, PPCs labelled with ‘Cell Trace Far Red DDAO-SE’ were counted from three 300 µm^2^ areas of flat mount images at each time point (**Table 2**). These cell counts showed significant improvement in retention of cells in the PKX-001-pre-treated groups at all three time points (P=0.001; N=3). As time progressed there were less cells at 6 months compared to at 3 months, suggesting some cells were dying.

**Table 2.** PPC cell counts in flat mount retina from each treatment group

| Time | Mutant<br>3 Months |  | Mutant<br>4.5 Months |  | Mutant<br>6 Months |  |
| --- | --- | --- | --- | --- | --- | --- |
| Treatment | Cells<br>only | Cells +<br>PKX-100 | Cells<br>only | Cells +<br>PKX-100 | Cells<br>only | Cells +<br>PKX-100 |
| Number of PPCs<br>per 300μm<sup>2</sup> | 739<br>±<br>2.90<br>(N=3) | 956<br>±<br>1.86<br>(N=3) | 684<br>±<br>2.90<br>(N=3) | 929<br>±<br>2.90<br>(N=3) | 510<br>±<br>1.96<br>(N=3) | 689<br>±<br>2.35<br>(N=3) |
| P value:<br>cells only versus<br>cells + PKX-100 |  | 0.001 |  | 0.001 |  | 0.001 |

### PPC integration and maturation in the host retina

To assess whether the PPCs had integrated into the host retina immunohistochemistry for the STEM121 marker was carried out. STEM121 is expressed in human cells from a variety of tissues including the central nervous system (including the eye), liver and pancreas. This antibody does not cross-react with rodent central nervous system tissue. Immunostaining for STEM121 was only seen in retinas treated with PPCs that were pre-incubated with PKX-001, and only at the 6 month time point. Immunostaining with STEM121 verified that “Cell Trace Far Red DDAO-SE”-positive PPCs were indeed human-derived PPCs (**Fig. 4**). At 6 months of age there remains only a single layer of host photoreceptor nuclei in the outer nuclear layer (ONL) and these did not label with STEM121. Some “Cell Trace Far Red DDAO-SE” staining however did not coincide with STEM121 staining. In such cases staining did not co-localize with definite cell bodies suggesting that this staining corresponded to PPCs degenerating and fragmenting with release of “Cell Trace Far Red DDAO-SE” into the extracellular space.

**Fig. 4.**
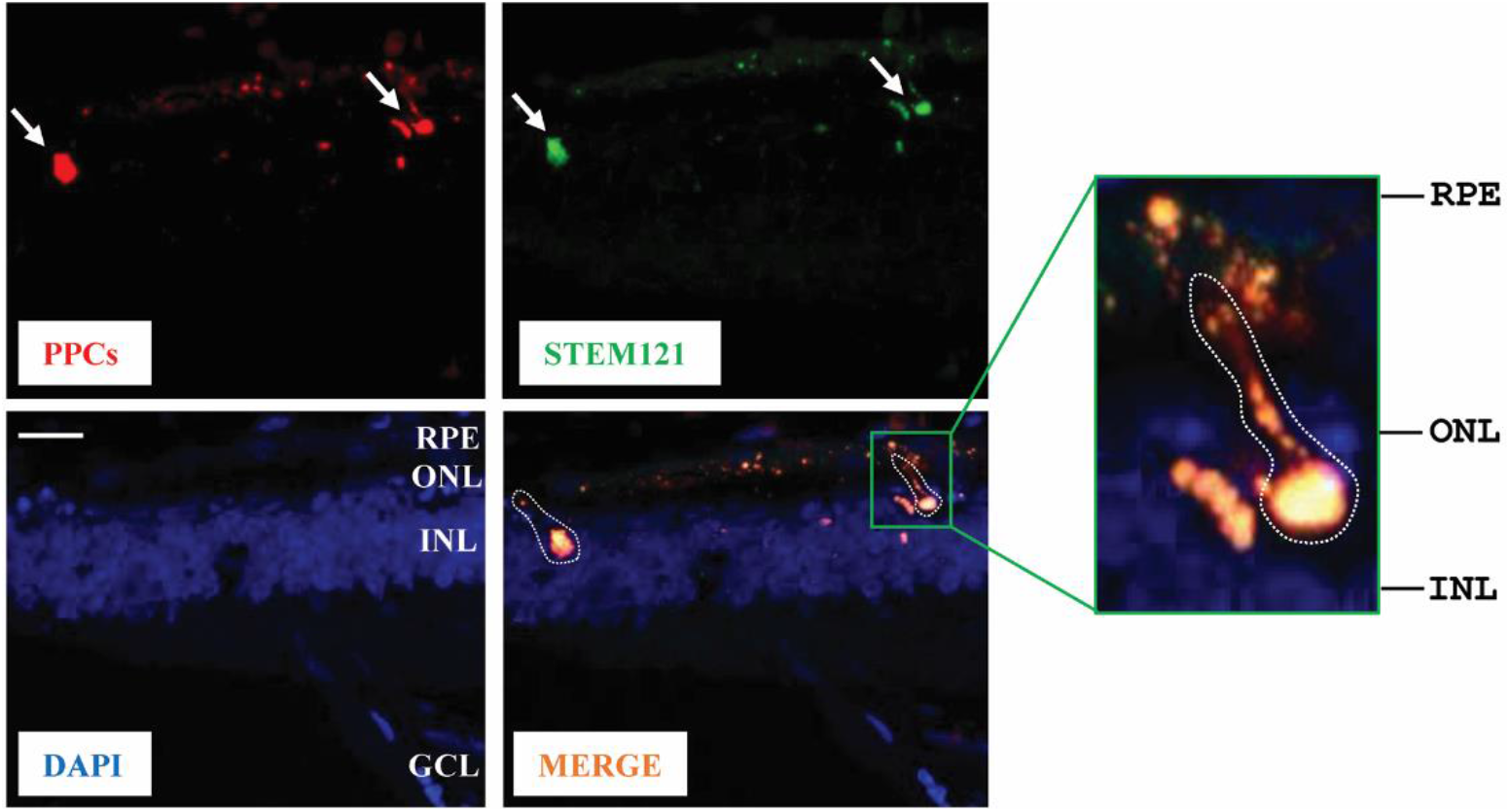
PPCs of human origin integrate into treated retina at 6 months. Representative sagittal cryosection through the retina showing PPCs labelled with Cell Trace Far Red DDAO-SE. The STEM121 immunolabeling in green identifies cells of human origin. Nuclei counterstained with DAPI. RPE, retinal pigment epithelium; ONL, outer nuclear layer; INL, inner nuclear layer; GCL, ganglion cell layer. Size bar = 25 µm. In the merged image two PPCs are identified by dotted lines. Inset shows higher magnification of one cell extending a projection towards the RPE.

Maturation of PPCs into retinal cells was determined by examining expression of the Far Red cell tracer and immunostaining with the human ribbon synapse marker Ribeye (C-terminal binding protein 2, CTBP-2), to identify synaptic connections between the host retina and the transplanted cells. We found examples of PPCs taking on an elongated profile reminiscent of mature photoreceptors (**Fig. 5A)** and expression of human Ribeye, suggesting that such cell maturation included making synaptic connections with host tissue (**Fig. 5B**). This staining was only seen at the 6 month time point where PPCs that were pre-incubated with PKX-001. No signs of maturation were observed in retinas treated with PPCs that had not been incubated with PKX-001.

**Fig. 5.**
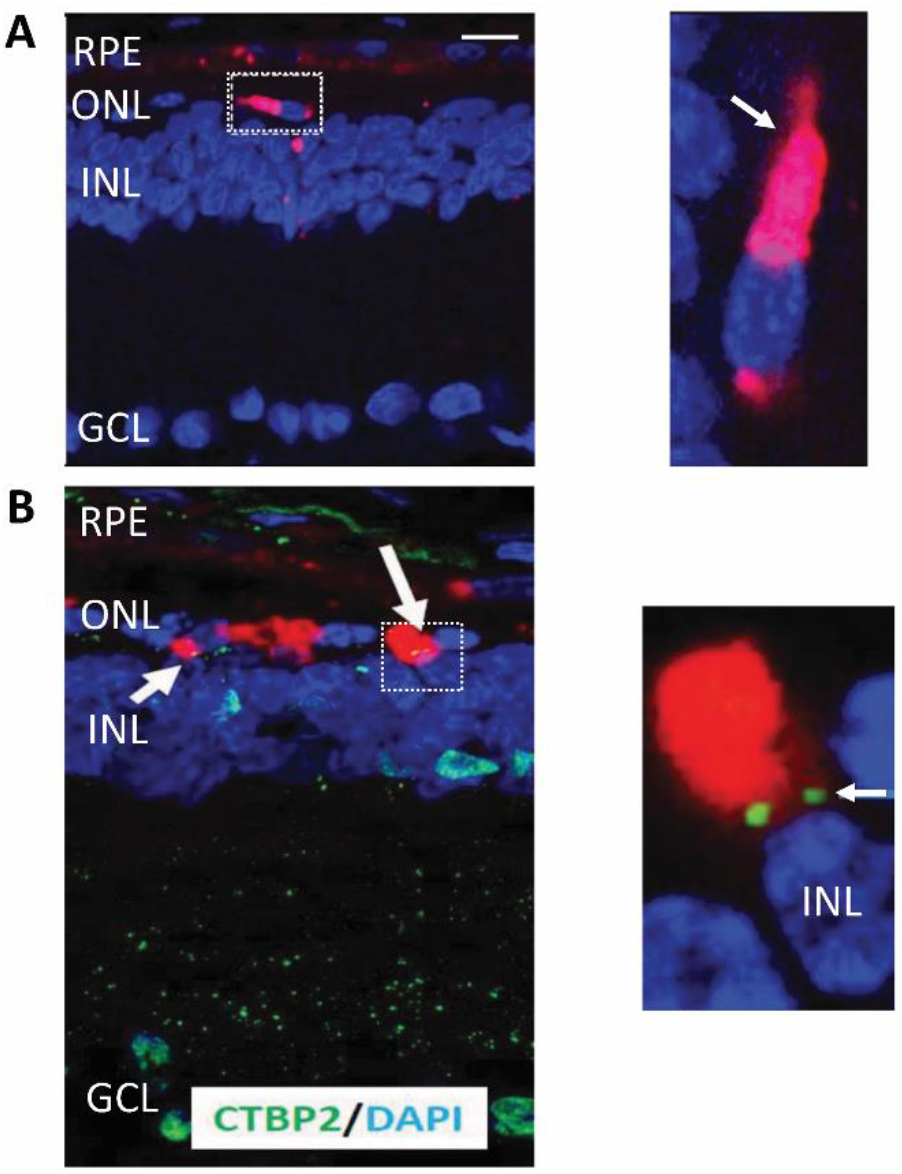
Maturation of PPCs in treated host retina at 6 months. (**A**) Expression of Cell Trace Far Red DDAO-SE in PPCs in the host retina. Nuclei stained with DAPI. RPE, retinal pigment epithelium; ONL, outer nuclear layer; INL, inner nuclear layer; GCL, ganglion cell layer. Size bar = 25 µm. Inset, higher magnification of a maturing PPC that has become elongated suggestive of and emerging inner segment (arrow). (**B**) Expression of Ribeye (CTBP2) in PPCs (arrows). Inset, higher magnification showing two characteristic green dots identifying the connection between the PPC and a second order INL neuron.

Finally, we examined the treated retinas for the expression of photosensitive opsin proteins characteristic of mature photoreceptors. We identified examples of fully differentiated cone photoreceptors with both an inner segment and an outer segment that expressed the photosensitive S-opsin protein (**Fig. 6A**). Similarly, we identified cone photoreceptors that expressed L/M-opsin in the outer segment process (**Fig. 6B**). Mature cells derived from PPCs were only found in retinas where the PPCs had been pre-treated with PKX-100 prior to transplantation at the 6 month time point. We did not detect any innate rat photoreceptors expressing these opsins as only a single row of nuclei in the ONL remained and these had all lost their outer segments.

**Fig. 6.**
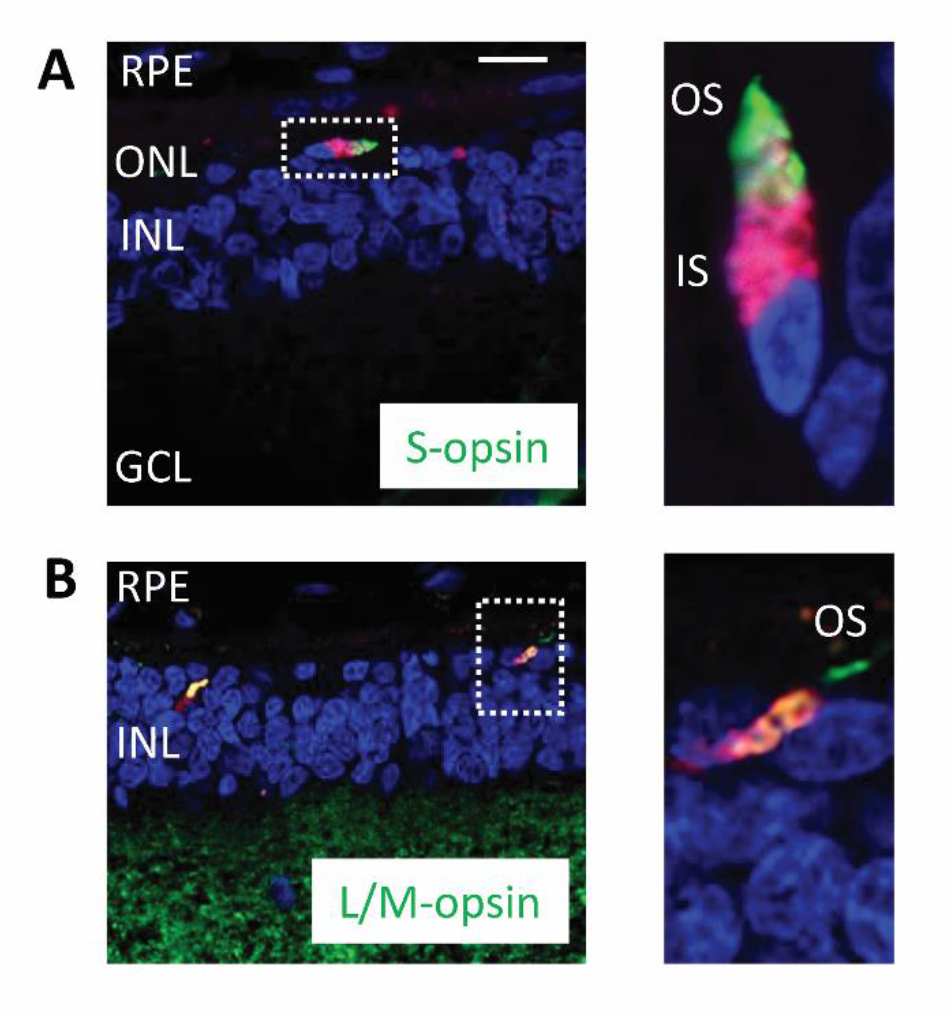
Expression of opsin proteins in treated retina at 6 months. (**A**) Confocal image of a cell expressing Cell Trace Far Red DDAO-SE (red) and immunolabeling for S-opsin (green). Nuclei stained with DAPI. RPE, retinal pigment epithelium; ONL, outer nuclear layer; INL, inner nuclear layer; GCL, ganglion cell layer. Size bar = 25 µm. Inset, higher magnification of a mature cone photoreceptor that has an inner segment (IS) labeled with Far Red and localization of S-opsin in the photoreceptor outer segment (OS). (**B**) Confocal image of cells expressing Cell Trace Far Red DDAO-SE (red) and immunolabeling for L/M-opsin (green). Inset, higher magnification showing a mature photoreceptor that has an inner segment (IS) labeled with Far Red and the outer segment (OS) localization of L/M-opsin.

## DISCUSSION

PKX-001 is an anti-aging glycopeptide. It is a small (544.2 Daltons), synthetic analog of the larger (>2,600 Daltons) family of compounds called Anti-Freeze Glycoproteins (AFGPs). These proteins occur naturally in Arctic and Antarctic fish, and other, cold-climate, dwelling invertebrates to enable cell and tissue function at subzero temperatures (DeVries, 1983, Bang et al 2013). Research in other species have shown much promise for AFGPs in the preservation of biological materials, and organ transplantation (Amir et al, 2005; Matsumoto et al, 2006, Deller 2014). Large-scale research has been limited however due to AFGPs’ costly extraction from un-sustainable sources, instability and large molecular size.

PXK-001 has been investigated via *in vitro* cell culture and in animal models under a variety of conditions and showed significant cytoprotective properties. *In vitro* studies in various cell lines has demonstrated significantly increased cell survival when PKX-001 is added to culture media and cells then exposed to various stressors (US Patent Nos. US20090311203). Supplementation with PKX-001 during *in vitro* culture of isolated human and murine islets has been shown to improve potency of transplanted islets and attenuate long-term tacrolimus-induced graft dysfunction (Gala-Lopez et al, 2016).

Our previous study using *ex vivo* human retina has also suggested that PKX-001 can help with human cell survival (Yanai et al, 2015). In that study we developed a novel culture system with human retina sitting on a bed of human retinal pigment epithelium derived from human embryonic stem cells, thus, mimicking the subretinal space. PPCs were sandwiched between the neurosensory retinal explant and retinal pigment epithelium. When PPCs were pretreated for 24 hours with PKX-001 (4 mg/mL), an almost 3-fold increase in PPC viability was observed at the end of a 10-day culture period.

Our current study extends this previous *ex vivo* study. Pre-treating PPCs with PKX-001 before subretinal transplantation results in improvement in PPC survival for up to 6 months. Most significantly though, signs of PPC maturation: objective functional benefits (electrodiagnostics and optical coherence tomography) and immunohistochemistry suggests that PKX-001 significantly improve PPC maturation into functional photoreceptors in a retinal dystrophy animal model. Despite that both S-opsin and L/M opsin were expressed in pre-implantation PPC we only observed their expression *in vivo* at the 6 month point and only in PPCs pre-treated with PKX-001. In addition, STEM121, a marker of viable human cells was only observed at six months and only in PKX-001 pre-treated PPCs. We interpret this as a sign that on subretinal injection PPC basal metabolism might be significantly suppressed by the hostile host environment and that at the 6 month timepoint, PKX-001 has significantly aided recovery to allow for photoreceptor maturation.

Currently, *in vivo* studies of PPC transplantation have produced mixed results in animal studies (Pearson RA et al., 2012; Barber et al., 2013; Gonzalez-Cordero A et al., 2013), and very limited benefits in human studies (Jin ZB et al., 2019, Wang Y et al., 2020). There is therefore a pressing need to improve transplantation methodology. The problems can be subdivided into two broad areas, issues of quality of PPCs and the hostile environment of the degenerating retina (Ikelle L et al., 2020). The most successful strategies so far have been improving the quality of PPC manufacture (Hirami Y et al., 2009, Jin ZB et al., 2012, West EL et al., 2012) a focus on immunosuppression (Nazari, H et al., 2015; Zhu J et al., 2017) and improvements in surgical techniques (Wang Y et al., 2020). A number of other strategies are also emerging. Small molecules are in development to overcome cross-species cell contamination. (Osakada F et al., 2009). Also, siRNAs that disrupt outer limiting membrane integrity and chondroitinase that digest chondroitin sulfate proteoglycans deposited as a result of glial scarring have also been used to improve the degenerating retinal environment for PPC transplantation (Barber et al., 2013). PKX-001 now add to these methodologies. Our studies suggest that PKX-001 works by ameliorating the toxicity of degenerating retina through a reduction of prostaglandin E2, a mediator that has been shown to stimulate cell death in several model systems (Takadera et al., 2004; Ricciotti et al., 2011; Miyagishi et al., 2013; Yannai et al., 2015).

In conclusion, in this preliminary *in vivo* study, PKX-001 significantly improves survival of PPCs in an animal model of inherited retinal disease. This study however was undertaken in a relatively small number of animals and further studies are needed to better quantify the effectiveness of the small molecule. Studies in other animal models are also needed since it has been shown that the genotype of an animal model can significantly influence the success of stem cell integration (Barber et al., 2013). In addition, more studies are needed to determine the precise mechanism of action underpinning the effectiveness of PKX-001. It has been shown that in some cases, functional restoration after transplantation can be attributed to RNA or protein transfer between graft and host photoreceptors instead of transplanted photoreceptors migrating and integrating into the photoreceptor layer of recipients (Santos-Ferreira T et al., 2016, Pearson RA et al., 2016, Nickerson PEB et al., 2018). It needs to be determined if this impacts on the method of action of PKX-001.

## ACKNOWLEDGEMENTS

The authors acknowledge financial support from Protokinetics Inc. in order to undertake this study. We also acknowledge the help of Dr. Evelina Rubinchik PhD in writing the manuscript.

